# Dynamics of p53 DNA binding sites contributes to functional selectivity of p53-driven gene expression

**DOI:** 10.1101/2021.09.18.460898

**Authors:** Jessy Safieh, Ariel Chazan, Pratik Vyas, Hanna Saleem, Yael Danin-Poleg, Dina Ron, Tali E. Haran

## Abstract

The tumor suppressor protein p53 is situated in the midst of a complex network that is activated in response to cellular stress. An unresolved question is how p53 activates its myriad target genes in response to the severity of the stress signal and consequently coordinates the functional outcome in a timely manner. We have previously shown that DNA torsional flexibility distinguishes among p53 response elements (REs). Here we calculated the flexibility of over 200 p53 REs. By connecting functional pathways of p53-dependent genes to the calculated flexibility of their REs, we show that genes belonging to pathways activated rapidly upon stress (e.g., cell-cycle arrest, energy metabolism and innate immunity) contain REs that are significantly more flexible relative to REs of genes involved in pathways that need to be more strictly regulated or are activated later in the response to stress (e.g., intrinsic apoptosis and p53 negative regulation). The global structural properties of several p53 REs belonging to the different pathways were experimentally validated. Additionally, reporter gene expression driven by flexible p53 REs occurred at lower p53 levels and with faster rates than expression from rigid REs. Moreover, analysis of published endogenous mRNA levels of p53 target genes as a function of the flexibility of their REs support our hypothesis. Overall, we demonstrate that DNA flexibility of p53 REs contributes significantly to the timely expression of p53 target genes and thereby plays an important role in cell-faith decisions in the p53 circuity.

## Introduction

Gene regulation depends on the correct spatiotemporal arrangements of transcription factors and their specific DNA response elements (REs), which leads to a productive interaction between them. It is now commonly accepted that recognition of DNA by regulatory proteins occurs through forming hydrogen bonds from the protein side chains to the DNA bases (termed direct or base readout) ^1^, and is supplemented by hydrogen bonds to the DNA backbone (termed indirect readout) ^2^, and by recognition of structural and dynamical (flexibility) properties of the DNA double helix at the transcription factor REs (termed shape readout) ^3^.

The tumor suppressor protein p53 is part of a complex network that is activated in response to various cellular stress signals. Activated p53 functions mainly as a transcription factor, regulating the expression of numerous genes, involved not only in various cellular pathways critical for preventing cancer, but also in pathways unrelated to cancer surveillance ^4, 5^. p53 is currently viewed more as the “guardian of homeostasis”, rather than the “guardian of the genome” ^6^, because it is currently appreciated that p53 responds both to acute DNA damage and to more “routine” activities, such as energy metabolism ^7-10^ and embryonic development ^11^. p53 in normal unstressed cells is present at low levels, caused by the association of p53 with the MDM2 protein ^12^. Upon exposure of cells to stress signals, cellular levels of p53 increase due to disruption of the p53-MDM2 interaction ^12^.

p53 has a multi-domain modular structure ^13, 14^. The N-terminus (NTD) contains a transactivation domain, the core domain contains a sequence-specific DNA binding domain (DBD), and the C-terminal domain (CTD) includes a tetramerization domain (TD) and a basic regulatory domain. The DBD of p53 binds to REs that consists of two decameric repeats of the form RRRCWWGYYY (R=A, G; W=A, T; Y=C, T), separated by 0-18 base pairs (bp) ^15, 16^. Base-pairs separating p53 half-sites (“spacer sequences”) are present in ~50% of all validated p53 REs ^15–17^. Full-length p53, as well as shorter protein constructs that incorporate the DBD, bind to their DNA targets cooperatively as tetramers, composed of a dimer of dimers ^13, 18-20^.

Precise discrimination of p53 multitude target genes is crucial for p53 to trigger correct cell-fate decisions. It is now clear that p53 responses to cellular stress signals are quite heterogenous, and vary with respect to p53 levels needed for target-gene activation ^21–23^ and the time elapsed since p53 induction (early versus late response) ^22^. However, the molecular mechanisms of p53 timely responses remain unclear. Two alternative models were proposed to explain differential gene expression upon p53 induction ^24, 25^. *The selective binding model* focuses on direct binding of p53 to its REs as the fundamental step of gene regulation by p53, and inherent differences in p53 REs as an essential driving force for discrimination between p53 target genes. *The selective context model*, suggests that p53 is not the crucial force behind target gene selectivity, but that discrimination is achieved by other events at the cellular and genomic levels. Within the selective binding model, base readout as well as indirect and shape readouts were all shown to contribute to the interaction of p53 with its target promoters ^26–34^. In particular, we have previously demonstrated experimentally that DNA torsional (twist) flexibility changes significantly between p53 REs that differ in the uncontacted WW (W = A, T) motif at the center of each p53 half-site ^27^, or at the RRR (YYY) bases abutting the central motif from either side ^34^. Moreover, we have shown that when p53 levels are low, the torsional flexibility of p53 REs positively correlates with transactivation levels, whereas when p53 levels are highly induced, transactivation levels positively correlate with p53 binding affinity ^28^.

The energy parameter deformability (V(B) in units of °^3^Å^3^) is a quantitative measure for the overall flexibility of B-DNA base-pair steps, that is their dynamic ability to deform (flex) under an external force, such as that arising from the binding of protein molecules ^35, 36^. We have previously determined that this rigorous measure of helical flexibility, that is dependent on all DNA helical degrees of freedom, has a strong and significant correlation with the torsional (twist) flexibility of p53 natural and consensus-like REs measured using cyclization kinetics experiments ^34^, and can thus be used to estimate the flexibility properties of p53 REs.

Here we demonstrate that upon classifying p53 target genes by the flexibility properties of their REs, we obtain a range of REs’ flexibilities – from very flexible to very rigid REs. Moreover, we show that genes that are expressed immediately upon p53 induction (when p53 protein levels are low) have flexible REs. On the other hand, we find that genes that are activated only later in the response to stress (when p53 levels are high) harbor very rigid REs. Using combined structural and functional analyses, in vitro and in cells, we establish here that DNA torsional flexibility of p53 REs has an important role in singling out the appropriate target sites for timely activation among the numerous p53-dependent promoters.

## Results and Discussion

### Classification of p53 REs by the functional categories of the nearby gene

For this classification we analyzed a set of 235 genes, previously studied by us ^34^, that abide by the following criterions: (a) the genes are directly activated by p53; (b) the REs of these genes lack spacer sequences between half sites; (c) their REs are located within the promoter of the first intron/exon of the gene. The specific function of each gene, the biological process in which it participates, and the consequent cellular outcome are summarized in Table S1. For statistical analysis of possible relationships between functional outcome and DNA flexibility these genes were organized into supergroups based on the cellular outcome of their action (Table 1). For example, cell migration involves cytoskeleton rearrangements, ECM degradation, cell shape changes etc. Moreover, these same processes are also part of axon guidance and wound healing. We grouped all these biological processes into a supergroup termed “Cytoskeleton/cell migration”. This functional categorization was applicable to 208 out of the 211 genes for which a p53-dependent function is currently known. The remaining three genes (PRDM1, PTP4A1, XPC), have two main functions one of which is p53 negative regulation that is common to all three genes ^37–39^ (Table S1 and Table S2). In this p53-centric study, we chose to assign the above three genes to the p53-negative-regulation group.

**Table 1.** End functional-outcome supergroups with at least five genes each (related to Figure 1)

| **Supergroup** | **Functional outcomes included in the group** |
| --- | --- |
| Cell adhesion/anti-cell-motility | Increase in cell-matrix adhesion; increase in cell-cell adhesion; anti-metastasis; anti-migration; anti-invasion; anti-angiogenesis |
| Cell growth | Cell division machinery; mitotic cell cycle; mitotic spindle organization; regulation of cytokinesis; symmetric cell division; cell proliferation |
| Cellular stress response | Response to: ER stress; metabolic stress, hypoxia, heat stress, cold stress; osmotic stress; unfolded protein response; ROS; oxidative stress; oxidative DNA damage; stress granules formation |
| Cytoskeleton/cell motility | Cytoskeleton organization; actin polymerization; ECM organization and degradation; muscle contraction; neuronal migration; movement along microtubules; directional migration; apical-basal polarity; cytoskeleton re-organization; axon guidance; migration; invasion; wound healing; EMT; platelet aggregation; angiogenesis; decrease in cell adhesion |
| Development | Embryonic development; morphogenesis; neurogenesis; synaptogenesis; muscle development |
| DNA repair | Damage repair; translesion DNA synthesis; synthesis of dNTPs |
| Early DNA damage response (DDR) steps | DNA damage response activation; cell-cycle arrest (G1/S and G2/M); p53 positive regulation; other events closely following DNA damage; immediate early genes for DDR |
| Energy metabolism | Energy metabolism; glucose metabolism; oxidative phosphorylation; ATP synthesis |
| Lipid metabolism | Lipid metabolism; lipid droplets; cholesterol metabolism; steroids metabolism; steroidogenesis |
| Extrinsic apoptosis | Death receptor pathway |
| Innate immunity | Innate immune response; inflammation; pyroptosis |
| Intrinsic apoptosis | Intrinsic apoptosis machinery and effectors |
| Ion homeostasis | Ion channels; ion transport; transport of small molecules |
| Membrane trafficking | Endocytosis; exocytosis; vesicle transport; secretory pathways; Golgi; membrane transport; vesicular lipid transport |
| p53 negative regulation | Targeting p53 for destruction; p53 down regulation after damage repair |
| Cell signaling | Signal transduction by various pathways, when the signaling event is followed by multiple possible functional-outcome results |
| Stem cells and differentiation | Stem cell biology; stem cells self-renewal; pluripotency maintenance and regulation; stem cells differentiation |
| Survival | Anti-apoptosis; anti-necrosis; anti-pyroptosis; anti other kinds of deaths; cell survival |

The set of 211 p53 target genes analyzed here harbor 228 20-bp binding sites. This is due to the observation that p53 target sites frequently contain more than two decameric repeats, and instead contain clusters of decamers, either abutting or distantly spaced. Cluster sites are known to confer additional binding affinity and increase transactivation levels from the nearby gene ^15, 40^. In order not to bias the analyses of the relationship between DNA flexibility and functional outcome from p53-dependent gene activation, by multiple REs of the same p53 target gene, we took only one 20-bp RE per gene for structural and sequence analyses. This was done both when the cluster sites were separated by a long intervening region (13 sites, Table S1), and when the sites were composed of more than two abutting decamers (seven sites, Table S1). In both cases we chose for analyses the more flexible full-site RE, based on the assumption that the more flexible RE is the primary and dominant RE of the cluster ^34^.

### Functional outcome categories of p53 activated genes have REs with a distinct structural signature

We calculated the DNA flexibility of REs belonging to the assembled set of the 211 p53-dependent genes, using the energy parameter deformability V(B) ^34, 36^. Next, we analyzed the relationship between the calculated REs deformability and functional outcome groups that contain at least five genes each, to minimize noise from small groups (total 18 groups, 187 genes). The variation in REs deformability between functional-outcome groups shows significant differences in their mean deformability values (*F* = 19.71, *P* = <0.0001, Figure 1, Table S3). The results show that p53 target genes that have REs with deformability values that are significantly above the mean are those that are needed for establishing a fast and timely p53-dependent response to cellular needs or acute stress: early DNA damage response steps, innate immunity, development, and energy metabolism (colored blue in Figure 1 and termed collectively as the “flexible REs category”). In contrast, p53 target genes that have REs with deformability values significantly below the mean belong to target genes that need to be strictly regulated, or that contribute to a later stage in the response to stress: intrinsic apoptosis, p53 negative regulation, cellular stress response and cytoskeleton/cell motility (colored red in Figure 1 and termed collectively as the “rigid REs category”). A post-hoc test established that each group within the flexible REs category is significantly (*P* < 0.0063) and solely different from each group within the rigid REs category but not from most of the remaining groups (Table S3). The largest difference is between development and cellular stress response (*Z* = 6.385, *P* <0.0001). Of note to p53 functions as a tumor suppressor protein, are the significant differences between early DNA damage response (DDR) steps and cellular stress response (*Z* = 5.03, *P* <0.0001), early DDR steps and intrinsic apoptosis (*Z* = 4.37, *P* =0.0019), and early DDR steps and p53 negative regulation (*Z* = 4.10, *P* = 0.0063). Concerning the groups with median flexibility, one can rationalize the mean deformability of several groups but not all. Lipid metabolism, like energy metabolism, is an early event, though it does not have as extremely flexible REs as does the energy metabolism group. The extrinsic (death receptor) apoptosis pathway has above-average mean deformability, although it is a death-related functional-outcome group, and hence expected by the our analysis to have deformability values that are below the mean. This may be related to the observations that extrinsic apoptosis is initiated by signals originating from outside the cell, such as natural-killer lymphocytes, or cytotoxic-T lymphocytes, and thus is intimately connected with the innate- and adaptive-immune systems ^41^, that by our analysis have above mean deformability values. A third example is the transition between cell-cycle arrest, DNA repair, and apoptosis that is known to be gradual and to depend on the accumulated level of DNA damage ^42^. Moreover, the regulation of DNA repair should be stringent, since erroneous DNA repair can lead to mutations and other chromosomal defects leading to cancer development ^43^. Thus, we expect DNA repair to have mean deformability between those of early DDR steps and intrinsic apoptosis, as can be observed from Figure 1.

**Figure 1.**
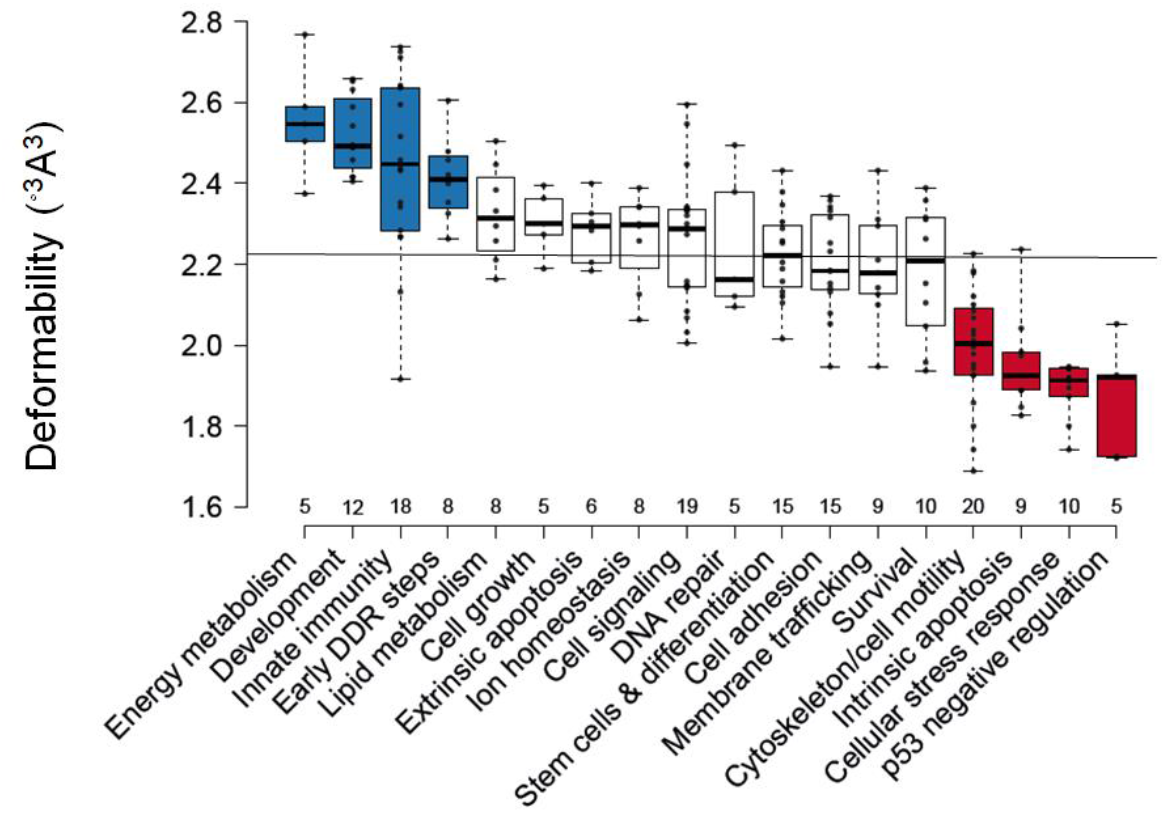
p53 REs belonging to functional-outcome groups that are needed early in the response to stress are more deformable than those needed later in the response to stress. Shown are functional-outcome groups with at least five genes each. All data points are shown as dots. The line across all groups marks the mean of means of the deformability of all analyzed groups. Center lines show the medians of each group. The figure was created by BoxPlotR ^72^. Box limits indicate the 25th and 75th percentiles, as determined by the R software. Whiskers extend to maximum and minimum values. Boxes are arranged by decreasing group average values from left to right. Blue boxes denote the “flexible REs category”, and red boxes signify the “rigid REs category”. For a list of group average values see Table S2, and for significant differences between the groups see Table S3. DDR = DNA damage response.

Interestingly, there were no difference in classifications to categories, between genes that have one full site REs and those that constitute a cluster of REs having two or more full site REs. In the flexible category there are four genes with clusters of REs (BTG2, p21, COL18A1 and PANK1), in the rigid category there are also four genes with REs clusters (BAX, MDM2, PMAIP1 and RFX7), and in the interim category there are five such genes (FHL2, GPC1, LIMK2, TGFA and TNFAIP8). Finally, the analysis was unaltered when excluding all sites containing RE clusters.

### Functional outcome categories of p53-activated genes have REs with a unique sequence signature

Sequence logos are a convenient way to depict base-pair preferences and information content of DNA binding sites ^44–46^. The sequence logo of the whole set of 211 REs of p53-dependent genes with known functional outcome (Figure 2A) shows a degenerate sequence pattern, similar to the pattern from previous studies ^15, 16, 34, 47^. Separating this overall sequence pattern to that of the two extreme categories (Figure 2B-C), one can observe increase and decrease in nucleotide preferences in various positions. To better evaluate quantitatively the differences in sequence preferences, we calculated the information content per nucleotide position (*Iseqs_l_*), of the set of aligned REs sequences per category (Figure 2D, see Materials and Methods for equations). Information content is related to the thermodynamics of protein-DNA interactions because it is a measure of the discrimination, by a regulatory protein, between binding to a specific binding site versus binding to a random DNA sequence ^48–51^. Comparison of the two extreme-functional-outcome categories (Figure 2D) shows that in the 5’ CWW position and the 3’ WWG position (CWW on the opposite strand) of the flexible REs category have higher information content (i.e., the sequence space is more constrained and thus sequences vary less within this category in these positions) relative to the rigid REs category (~47% higher, respectively). On the other hand, the category with rigid REs is more constrained at the 9^th^ nucleotide, in the middle of the inner YYY tract, and in the 11^th^ position. The inner YYYRRR region of the rigid REs category has 33% higher information content than that of the flexible REs category.

**Figure 2.**
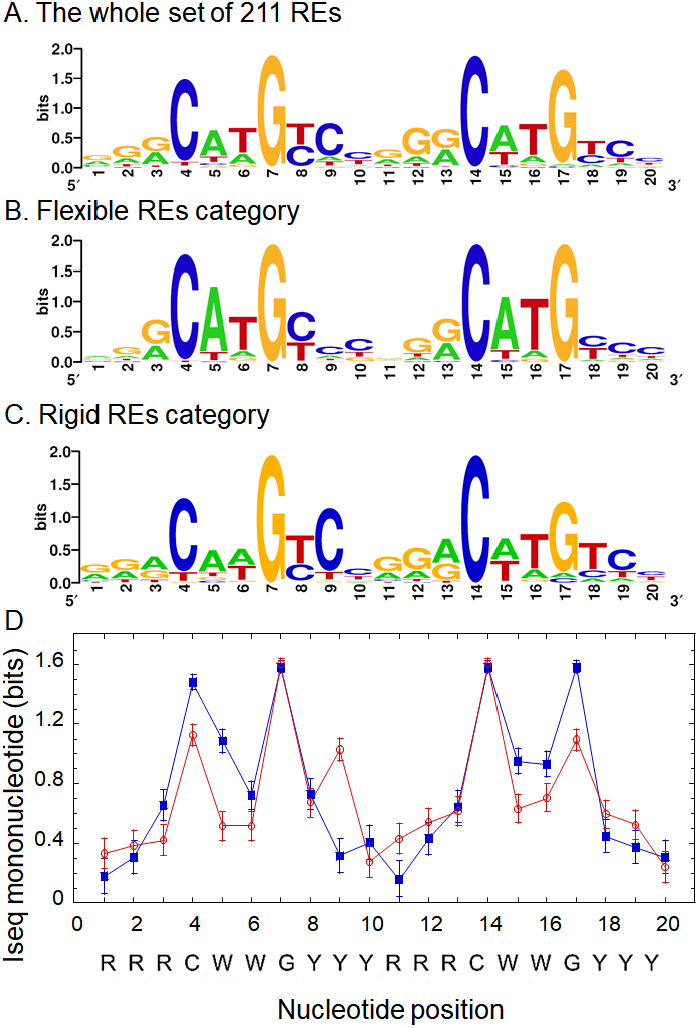
Sequence logos and information content of the studied REs. (A) All 211 p53 REs without spacers and with known functional outcome of p53 activation studied here. (B) REs belonging to the flexible category. (C) REs belonging to the rigid category. All sequences were aligned by transcription direction before generating the sequence logos. Sequence logos were generated by weblogo ^45^. (D) Information content per position (Iseql) on the mononucleotide level for the REs belonging to the flexible category (blue squares) versus those belonging to the rigid category (red circles).

The observation of high information content at the two CWWG motifs of the flexible REs category is due to the almost exclusive preference for the flexible CATG sequences in both half-sites. p53 does not directly contact the WW doublet in the complex with its REs ^26, 29-31, 33, 52, 53^, and thus DNA recognition in this region is by shape readout. We have previously shown that CATG-containing p53 sites are more torsionally flexible than CAAG and CTAG containing sites ^27^. This is due to the flexible CA and TG dinucleotides making this motif. The high flexibility of CATG-containing REs is the reason for the high kinetic stability of p53 dimers on these sites, which facilitates the swift assembly of functional p53 tetramers from DNA-bound p53 dimers ^28^. The CAAG and the CTAG motifs are rigid, and hence require higher p53 levels for forming functional tetramers, as the dimeric species are not stably bound to such sites ^28^. However, it is likely that this destabilization of the CAAG and CTAG motifs is partly compensated for by the stabilization of the p53-DNA interface via hydrogen bonds between Lys120 and the central G base in the opposite strand to the internal YYY tract ^26, 29-31, 33, 52, 53^. This may explain the stringent requirement for a C9 in the middle of the internal YYY tract in the rigid REs category. Thus, our results suggest that various forces optimize p53-REs interactions selectively. In the flexible REs category, high information content (binding free-energy contributions of each base) in shape-readout positions comes together with high deformability (structural property of regions within the DNA double helix). These features of the DNA double helix within the flexible REs category all contribute to a possibility for a fast response upon stress. In the rigid REs category, a direct-readout contact (Lys120 to G base) compensates for low binding affinity in the shape-readout positions.

It has been suggested that the distinction between pro-survival and pro-apoptotic genes resides at least in part in the increased frequency of A/T versus G/C bps at positions 3, 8, 13, and 18 of p53 REs, respectively, regardless of the sequence of the CWWG core motif ^54^. This suggestion was based on the authors study of DNA binding by the Lys120Arg p53 mutant, and on an analysis of p53 REs of 16 pro-survival versus 16 pro-apoptotic genes ^54^. Indeed, it is known that when the sequence is G1G2A3, the binding affinity to p53 is higher than when the sequence is G_1_G_2_G_3_ or A_1_G_2_G_3_, where in the last two cases Lys120 binds to both G_2_ and G_3_ bases, and in the first case to G_2_ and the T base on the opposite strand to A_3_ ^26^. However, our results using 42 flexible REs pro-survival genes, and 47 rigid REs pro-death genes show that other signals are more important for the distinction between survival and death. First, results from this study, as well as several previous studies ^15, 27, 55^ strongly suggest that the central CWWG vary between pro-survival genes, such as cell-cycle arrest, where it is CATG, shown to be more torsionally flexible ^27^, to pro-death genes, such as intrinsic apoptosis, where it is CAAG, shown to be more torsionally rigid ^27^. Here we add that a highly conserve C base at position 9 is important for optimal direct contact of Lys120 to the G base on the opposite strand, especially in the rigid REs category, where the contact to the central CAAG motif is sub-optimal.

### Experimental validation of the flexibility of five natural p53 REs by cyclization kinetics

Cyclization kinetics of DNA minicircles is a robust method to measure the global structural and dynamical properties of DNA molecules ^56, 57^, and it is carried out by following the rate of ligase-catalyzed closure of DNA molecules with cohesive ends into small rings. The REs chosen for the current study belong to genes that function at different cellular pathways (Table 2), and they were chosen by their deformability values (Table 2, rightmost column). As in our previous studies on p53 binding sites ^27, 34, 58^, the only significant change in global-structural parameters of these sites is in the twist (torsional) flexibility of the DNA binding sites (Table 2, Figure S1). Moreover, the results of the experimentally-derived torsional flexibility values showed similar trend to the values of the calculated deformability (Table 2). Thus, rigorous experimentally-derived torsional-flexibility values, show that CCNG1, RRM2B, and p21-5’ are the most flexible sites in this set, whereas PMAIP1 and TP53AIP1 are the most rigid sites. This experimentally corroborates our hypothesis that p53 REs that belong to the early DDR steps group are more flexible than those that belong to the intrinsic apoptosis pathway.

**Table 2.**
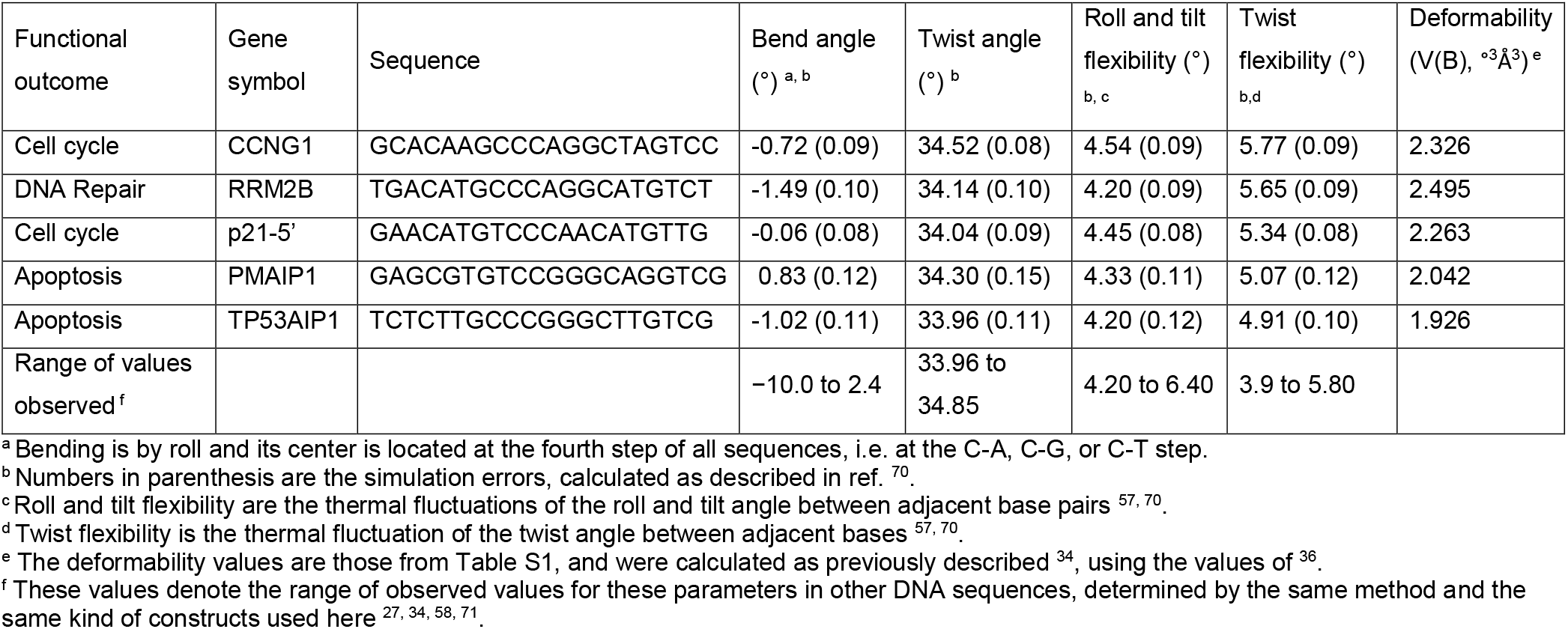
Best-fit parameters from cyclization kinetics measurements for the sequences studied here

### p53 target gene activation as a function of p53 levels

To further validate our hypothesis in a cellular context we carried out luciferase reporter gene assays in human non-small lung carcinoma cell line (H1299), ectopically expressing p53 along with reporter plasmids, each carrying a distinct RE. First, we ascertained that p53 ectopic expression is linearly correlated with the amount of transfected p53 expression plasmid pC53-SN3. This was done by transfecting cells with increasing amounts of p53 expression vector and immunoblotting the cell lysate (Figure S2A). Densitometry analysis of the observed p53 band, normalized to the β-actin band, confirmed the linear relationship between p53 protein level and pC53-SN3 amount (Figure S2B, R = 0.998). We then carried out reporter-gene assays for the five p53 binding sites described above at five different p53 levels (Figure 3). We analyzed transactivation measured by luciferase activity 48 h after transfection, to enable observing a significant signal also in cells transfected with the lowest p53 plasmid amount (1 ng p53 plasmid). Results show that transactivation rises as a function of p53 levels (Figure 3). One-way ANOVA, followed by mean comparisons, revealed that transactivation levels of the five genes are significantly different from each other at 1, 2, 5,10, 25ng of pC53-SN3 plasmid (Figure S3A-E, *P* < 0.005), but are similar at 75 ng of pC53-SN3 (Figure 2A, Figure S3F).

**Figure 3.**
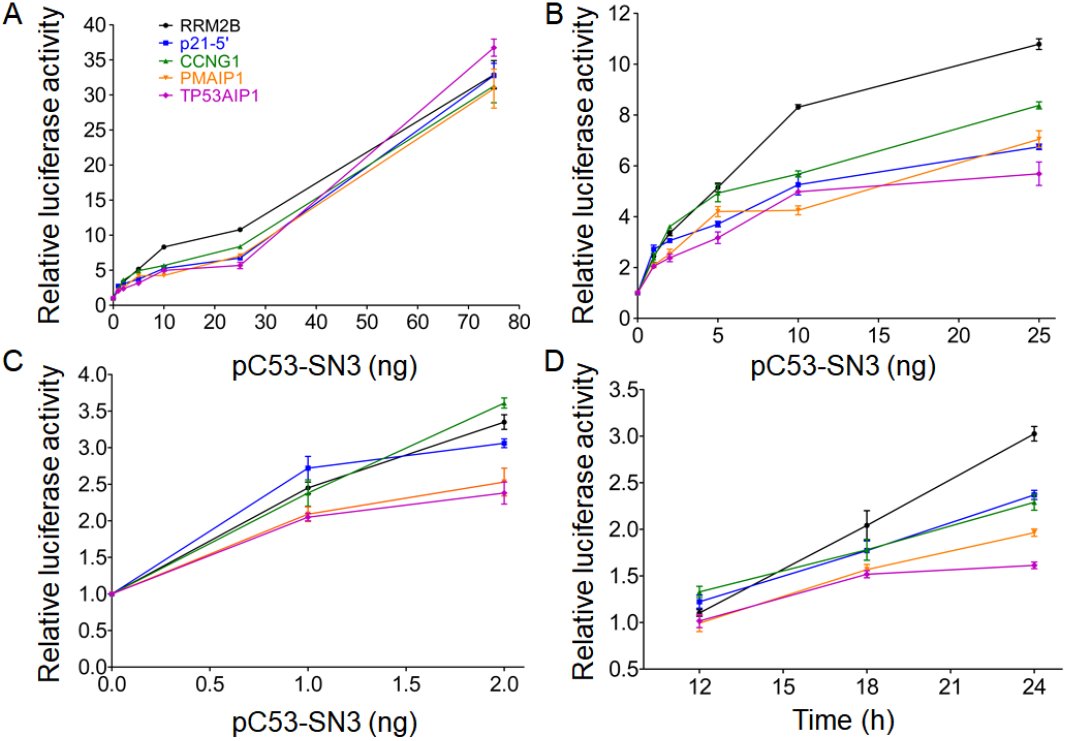
Transactivation from p53 REs as a function of p53 plasmid levels and of time. Fold increase in transactivation level was measured from three flexible p53 REs (CCNG1, RRM2B, and p21-5’) and two rigid REs (PMAIP1, and TP53AIP1) as a function of six different p53 expression levels, 48 h post-transfection (A-C), and additional three time points, using 25 ng p53 plasmid (D). Luminescence values were normalized to the transfection efficiency of co-transfected, constitutively expressed Gaussia luciferase (Gluc). Results were further normalized first to the empty Cypridina secreted luciferase (CLuc) vector, and then to results obtained without p53. Error bars represent the mean ± SD of four to eight independent experiments, each containing two technical replicas. (A) The whole range, 1-75 ng p53 plasmid. (B)-(C) for clarity these panels show the 1-25 ng and 1-2 ng range, respectively.

Focusing on cells expressing very low levels of p53 (1 ng p53 plasmid, Figure 3C), results show that transactivation from two flexible REs (RRM2B, and p21-5’) was distinctly higher compared to transactivation from the two rigid REs (PMAIP1 and TP53AIP1, *F* =4.96, *P* = 0.0044, Figure S3A). When 2 ng p53 plasmid were used (Figure 3C), transactivation levels from each of the three flexible REs were significantly higher than that from the two rigid REs (*F* = 22.9, *P* < 0.0001, Figure S3B). At a higher p53 plasmid level the PMAIP1 and p21-5’ REs are not monotonously increasing in transactivation levels, as the other REs are. Thus, individual pair-wise differences between REs are significantly different only when comparing the two most flexible REs - CCNG1 and RRM2B, to the two rigid REs – PMAIP1 and TP53AIP1 (Figure 3B, Figure S3 C-E). The transactivation from cells transfected with all tested p53 plasmid amounts, except 75 ng, was correlated with the experimental torsional (twist) flexibility of the five studied REs (Pearson *r (P*) = 0.35 (0.03), 0.74 (<0.0001), 0.64 (0.004), 0.84 (<0.0001) at 1, 5, 10, and 25 ng p53 plasmid, respectively), being most strongly at 2 ng plasmid (Pearson *r* = 0.91, *P* < 0.0001). This is consistent with our previous results ^28^, that at basal levels of p53 that is a strong and significant correlation between transactivation and DNA flexibility of p53 REs.

We compared known amounts of recombinant full-length p53 protein to transfected levels of p53 plasmid in H1299 cells (Figure 4). In cells harboring 1 and 2 ng p53 plasmid, we estimated that p53 protein levels are approximately 3.7 X 10^-4^ and 7.7 X 10^-4^ pmole/μg of whole-cell extract (WCE), respectively, which is similar to basal p53 levels previously found in HCT116 cells ~5 X 10-4 pmole/microg WCE ^59^. At 75 ng p53 plasmid there are 3.6 X 10^-3^ pmole/μg WCE, which is similar to endogenous p53 levels found after stress induced by camptothecin treatment in HCT116 cells 11 X 10-4 pmole/microg WCE ^59^. Thus, the results ascertain that the amount of p53 plasmid used in this transiently-transfected plasmid study corresponds to known levels of endogenous p53 protein that range from basal levels to those induced by cellular stress.

**Figure 4.**
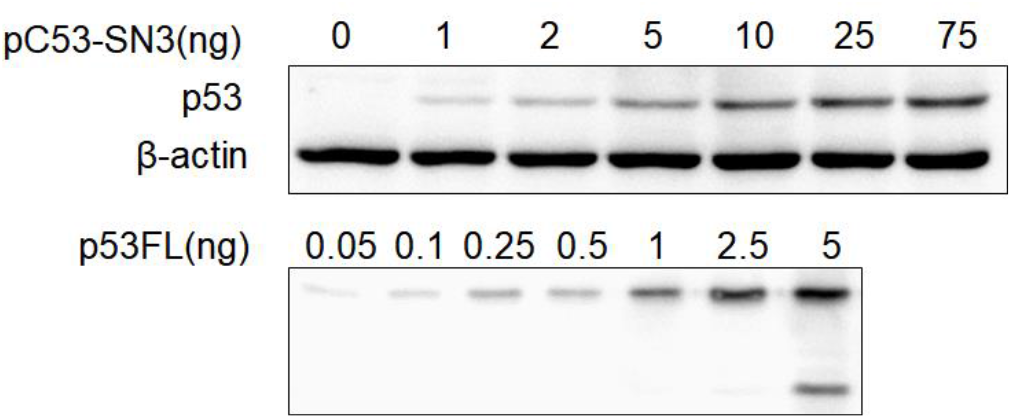
Estimation of the amount of p53 protein in H1299 cells upon transfection with various amounts of p53 plasmid. Upper panel shows H1299 cells transfected with increasing amounts of p53 expression vector, as described in Figure S2. Lower panel represents a serial dilution of purified full-length p53 recombinant protein. The lower band in the 5-ng lane is a degradation product, which was included in the quantification.

### p53 target gene activation as a function of time

The higher transactivation levels observed 48 h post-transfection from flexible REs, relative to rigid REs, could be due to an earlier start of transactivation, or alternatively due to faster transactivation rates. To distinguish between these possibilities we examined transactivation levels, at 25 ng p53 plasmid (to enable observation of a significant signal also at shorter times), at additional time points (12, 18, and 24 h post-transfection, Figure 3D). One-way ANOVA analysis at 12 h post-transfection revealed a significant difference between the transactivation level of the five REs (*F* = 4.34, *P* = 0.008), which remains significant at 18 h (*F* = 4.36, *P* = 0.008) and 24 h (*F* = 79, *P* < 0.0001). A post-hoc analysis indicated that at 12 h post-transfection, transactivation from CCNG1 and p21-5’ was significantly different from all rigid REs (Figure S4A). At 18 h post-transfection, post-hoc analysis indicated that transactivation from RRM2B was significantly different from all rigid REs (Figure S4B). At 24 h post-transfection, transactivation from all flexible REs studied here was higher than transactivation by all rigid REs (Figure S4C). Even though p53 levels in our study are due to ectopic expression of p53, the time frame for observation of post-transfection transactivation activity are in line with studies probing p53 activation due to induction of cellular stress events ^60, 61^. A positive significant correlation is observed between p53-dependent transactivation level from the five studied REs, at all studied post-transfection time points, and their experimental twist flexibility (*r* = 0.50, 0.53, and 0.77, and *P* = 0.004, 0.001 and *P* < 0.0001, for 12, 18, and 24 h, respectively).

### Hafner’s time-dependent RNA-seq study of endogenous mRNA levels of p53 target genes support the division to functional outcome categories based on extreme REs flexibility

To further support our results and to be able to mimic a more natural environment, where transcription is affected by multiple other factors, we analyzed the time-dependent RNA-seq data from the study by Hafner et al. ^62^. Hafner et al. studied the relationship between p53 dynamics and p53-dependent gene expression following DNA damage caused by γ-irradiation at low p53 levels, which triggers a series of p53 pulses ^62^. The measurements were taken every hour between 1 h and 12 h, and another measurement at 24 h. Their results showed that the RNA-seq measurements can be clustered into five distinct clusters according to gene expression dynamics ^62^. We analyzed RNA-seq measurements of p53 target genes that belong to cluster 1 (the cluster with the most enriched p53 bound genes) and for genes that appear in our set (18 of 32 genes in cluster 1). Since the fold-change of mRNA expression (relative to the expression level at time zero) changes as a function of time, we normalized fold-change values to the average fold-change of all selected genes at a particular time point. Likewise, we used normalized deformability values (Table S1). We then plotted the normalized fold-change of gene expression as a function of normalized deformability values of each specific gene. Values are ranked relative to the average value of the parameter (i.e., the lines drawn across both axes, Figure 5). From Figure 5 it can be observed that early DDR steps genes (colored blue) have above average flexible REs, and are expressed at above average level (>2 normalized fold change) at every tested time point. On the other hand, genes belonging to the extremely rigid REs category (colored with various shades of red) are all below average rigidity and are expressed at below average level. The only exceptions are the two genes from the cellular stress response group, SESN1 and SESN2, as well as MDM2 from p53 negative regulation group. Nonetheless, these genes also express rather low levels of mRNA (<2 normalized fold change, Figure 5). MDM2 is known to have a second full site, 17-bp away from the first one, and as discussed above, additional p53 REs can lead to enhanced transactivation levels ^15, 40^. This could perhaps be the case also for SESN1 and SESN2, as additional p53 REs (or p53-like REs) clusters can be observed in the UCSC Genome Browser for these genes. Analysis of such small data set is problematic, yet, differences in expression levels were seen only between the flexible REs group (early DDR steps) and the rigid REs group (intrinsic apoptosis, cellular stress response, cytoskeleton, and p53 negative regulation), at all the studied time points (*P* < 0.004). All in all, analysis of the results of Hafner at al. ^62^ support our hypothesis that DNA flexibility is an effective indicator for the level of p53-activated gene expression at low levels of p53, and that genes that are expressed early after stress induction have flexible REs and are mostly expressed at higher levels than those that are expressed at a later stage in the response to stress. At present, no conclusions can be drawn for genes belonging to groups with intermediate flexibility REs with regard to their level of mRNA expression. Studies of a larger set of p53-dependent genes at high temporal and positional resolution will help to further substantiate these observations in natural cellular environment.

**Figure 5.**
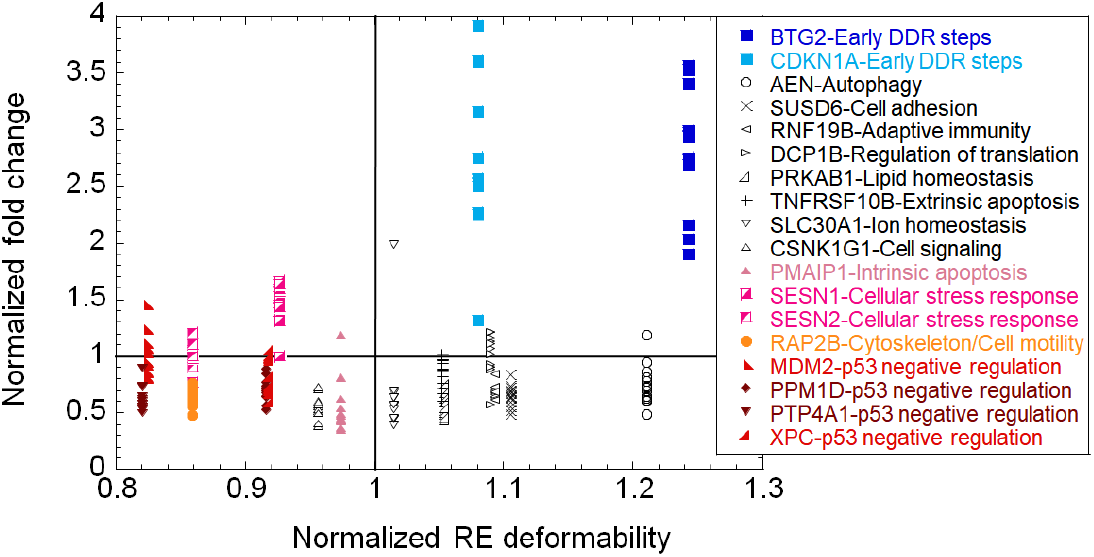
Endogenous mRNA levels from p53-dependent genes are grouped by their RE deformability. The data shown is based on the RNA-seq results from Hafner et al. ^62^. Fragments Per Kilobase Million (FPKM) values of each gene (i), at each time point (j) were normalized to their FPKM value at T = 0, arriving at fold-change values. Then these values were further normalized to the average fold-change of all genes at each particular time point (j). That is, each point is calculated from the equation: 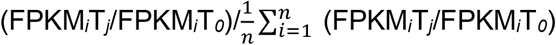. Deformability of genes’ REs was normalized in the same manner. Plotted are the normalized fold-change values as a function of normalized deformability values. The lines crossing each axis denote values that are above/below the average value of the parameter shown in each specific axis.

## Summary and Conclusions

Orchestrated expression of p53 target genes governs cell-fate decisions. An important question in the field is how p53 discriminates between the myriad p53-dependent target genes. The importance of differential binding affinities of p53 to its REs ^55^, the dependence of cell-fate decisions on p53 levels ^21–23^, and the elapsed time after p53 induction ^22^ have been widely acknowledged. However, the mechanistic basis for timely p53-dependent gene expression is currently unknown. In our previous studies, we showed that variation in torsional flexibility as a function of the RE sequence is the major structural feature that differentiates between p53 REs ^27, 34^. Moreover, at high p53 levels, transactivation from a given RE, is regulated by p53 binding affinity whereas at basal p53 levels it is regulated by p53 binding kinetics, governed by the DNA flexibility of the RE ^28^. These mechanistic observations support selectivity in p53 binding for understanding target-gene discrimination in the p53 system. Yet, it does not explain how p53 can coordinate its myriad target genes in an orderly fashion. Here we show that DNA flexibility is a key factor in determining p53 functional outcome. Transcription of genes required for fast response, at low p53 levels, is associated with REs characterized by high DNA flexibility. As we have shown previously ^28^, DNA torsional flexibility enables a stable binding of p53 dimers to p53-dependent REs also when p53 levels are low. Under these conditions, p53 tetramers can be formed sequentially (i.e., at low binding cooperativity) without the need for p53 accumulation over an extended time span. Therefore, our results, obtained in-vitro and in cells, indicate that the information for the selective binding of p53 to DNA is intrinsically encoded within the base sequences of the p53 REs. Hence, in addition to the statement that “p53 is smart” ^25^, the current data demonstrate that the DNA of p53 response elements is “smart” - the REs base sequences are information-rich by virtue of their structural properties. Consequently, the timely expression of genes’ activation is another fundamental code within the layers of codes that co-exist within the DNA double helix.

Our findings that REs flexibility contributes significantly to the orderly expression of p53-activated genes is in line with the observations that the presence of a single p53 full-site RE is sufficient for activating p53-dependent gene expression, without the need for cooperation from additional TFs ^63, 64^. This does not preclude combinatorial control by co-activating TFs at some genomic binding sites ^65^, as well as effects originating from chromatin context ^66, 67^, or from post-transcriptional events, such as mRNA stability of p53 target genes ^62^.

## Materials and Methods

### Functional annotation of p53 target genes

We used here a previously assembled set of 235 p53 target genes that are all upregulated by p53, are validated for binding in cellular context and for p53-dependent gene expression, and reside in the promoter of the first exon/intron region ^34^. This set of genes harbor 250 p53 REs that are all without spacer sequences between half sites, because p53 REs with long spacer sequences between half-sites (>9bp) can be bound by p53 in hemi-specific mode, a binding mode that nonetheless has a comparable binding affinity and transactivation levels to that of the full-specific complex ^58^. We modified the previous dataset by deleting the DDIT4 REs, because this gene was shown to be directly upregulated by RFX7 and not by p53 ^68, 69^ and added two genes RhoC and Scrib (Table S1). We located each gene in the Genecards suite (https://www.genecards.org/) by their Entrez ID, gene symbol, and gene name. We then looked for functional annotations of these genes in the Gene Ontology (GO) project (under the ‘Biological process’ and ‘Molecular function’ categories) through UniProtKB/Swiss-Prot (https://www.uniprot.org). GO annotations were supplemented by publications retrieved through individual literature searches for each gene product, and especially those centering on the role of these genes in the p53 system.

### Cells, plasmids, and reagents

H1299 human non-small lung carcinoma cells (p53-null cells) were obtained from ATTC (CRL-5803). They were thawed and propagated in RPMI-1640 (Biological industries, Beth Haemek, Cat # 01-101-1A) with 10% Fetal Bovine Serum (Biological industries, Beth Haemek, Cat # 04-127-1A), 2 mM L-glutamine, and 1% Pen-Strep solution (Biological Industries, Beth Haemek, Cat # 31-031-1C). All experiments were performed in this medium. Cells were maintained in a humidified incubator at 37°C and 5% CO_2_. Cells were routinely checked for negative mycoplasma infection.

The reporter gene vectors pCLuc Mini-TK 2, encoding Cypridina secreted luciferase under the control of a minimal promoter, and pCMV-GLuc, encoding constitutively expressed Gaussia secreted luciferase, were purchased from Thermo-Fisher. pC53-SN3 construct (where p53 expression is driven by the CMV promoter) was used for wild-type p53 expression in H1299 cells. p53 expression plasmid was kindly provided by Varda Rotter, Weizmann Institute of Science, Rehovot, Israel.

### Generation of reporter plasmids containing p53 REs

All DNA sequences were synthesized by Sigma Genosys (Israel), purified by standard desalting, and are described in Table S4. The sequences corresponding to p53 REs were incorporated in pCluc Mini-TK plasmids using site-directed mutagenesis with KAPA HiFi™ DNA Polymerase kit (Gamidor Diagnostics), and transformed into competent DH5α cells, according to manufacturer recommendations. Overexpressed plasmids were extracted using QIAprep^®^ Spin Miniprep Kit (Qiagen), according to the manufacturer’s protocol. The identity of incorporated sequences was verified using Sanger sequencing.

### Reporter gene assays

H1299 cells were seeded in 24-well plates at a density of 5X10^4^ cells/well and incubated for 24 h. Then cells were co-transfected with 800 ng of either pCLuc Mini-TK 2 harboring p53 RE or a corresponding empty vector along with 50 ng of pCMV-GLuc and 0 to 75 ng (0, 1, 2, 5, 10, 25, 75 ng) of p53 expression vector (pC53-SN3). The total DNA amount was adjusted using pcDNA 3.1 plasmid. Transfection was performed with jetPEI (Polyplus) reagent according to the manufacturer’s recommendation. Transfection efficiency was monitored by transfecting separate wells with 0.1 μg eGFP along with Gluc and pcDNA plasmids. An aliquot of growth medium (20 μl) was taken at various time points post-transfection and luminescence was measured using Pierce™ Cypridina Luciferase Flash Assay Kit and Pierce™ Gaussia Luciferase Flash Assay Kit (both from Thermo Fisher) according to the manufacturer’s protocol. Signal was detected by CLARIOstar 96 well plate reader (BMG LABTECH). Four to eight independent experiments were carried out, each containing two technical replicates.

### Analysis of transactivation from p53 REs

To correct for transfection variability, all results from Cypridina Luciferase luminescence were normalized to Gaussia Luciferase luminescence. Next, the results were normalized in two steps. First, the results obtained with pCLuc Mini-TK 2 harboring p53RE (Cluc^+RE^) were divided by the results obtained with the corresponding empty vector (Cluc^-RE^). Second, the results obtained with p53 (1-75ng of pC53-SN3) were divided by the results obtained without p53 (0 ng of pC53-SN3). Fold increase in transactivation is thus given by the equation:

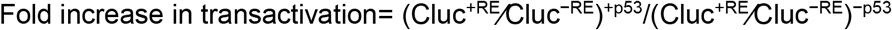

### p53 protein quantitation and detection

Cells were lysed in RIPA buffer (150 mM NaCl, 1% NP-40, 0.5% Na deoxycholate, 0.1% SDS, 25 mM Tris-Cl, pH 7.5) containing 1/100 (Vol/Vol) Protease inhibitor cocktail (Sigma, Cat # P8340). An equal amount of protein from the different samples were resolved on 10% SDS PAGE, and transferred to PVDF membrane. The blots were subsequently incubated overnight at 4°C with primary antibodies against p53 (DO-7 mouse mAb, Cell Signaling, Cat # 48818) and β actin (13E5 rabbit mAb, Cell Signaling, Cat # 4970). Membranes were then reacted with a secondary antibody (Anti-mouse IgG, and Anti-rabbit IgG conjugated to Horseradish Peroxidase (both from Cell signaling, Cat # 7076 and # 7074, respectively), developed using Western Bright^TM^ enhanced chemiluminescence reagent (Advansta), air-dried, and exposed using Fusion Pulse (Vilber) detection system, followed by quantitation using Evolution-Capt software. For p53 serial-dilution curve, full-length recombinant p53 protein was resolved on gels together with the transfection samples.

### Cyclization kinetics and simulation of cyclization data

DNA constructs for cyclization kinetics were synthesized using the top library/bottom test sequence PCR scheme, described previously ^27, 34, 57, 58^, except that here we used as test sequences DNA sequences containing the full 20-bp long p53 sites plus 5-bp flanking sequences on either side, specific for each natural p53 site (see Table S5 for sequences), because we have previously shown that sequences flanking p53 REs modulate p53 binding and transactivation ^34^. However, the global structural characteristics of the sites (Table 2, Figure S1) are those of the central 20-bp p53 sites without flanks, to concur with our previous studies ^27, 34, 58^. The DNA test sequences for cyclization experiments (bottom PCR templates) were synthesized by Sigma Genosys (Israel), whereas the library DNA sequences (top PCR templates) and the fluorescein- and tetramethylrhodamine (TAMRA)-labeled oligonucleotide primers were synthesized by the Keck foundation Laboratory at Yale University. Cyclization kinetics measurements were carried out as described previously ^27, 34, 58^. PCR reactions (50 μl) contained 1× PrimeSTAR buffer, 0.2 mM dNTP mix, 0.3 μM of each of the primers, 100 nM of the top and bottom templates, and 0.04 units of PrimeSTAR DNA polymerase (Takara, Japan, Cat # R010B). Ligase concentration was varied as a function of phasing length (0.08 U/μl for the in-phase 156L14 and 156L16 constructs and 1.0 U/μl for all other phasing constructs) and total length (from 0.08 U/μl for the 157 constructs, to 0.8 U/μl for the 154–155 and 158–160 constructs and 2.5 U/μl for the 150–153 constructs). We derived quantitative data on the conformational properties of the test sequences by simulating the cyclization data as previously described ^27, 34, 58, 70^, using the simulation program developed by Zhang and Crothers ^70^. The outcome of the simulations are the bend angle, twist angle, roll and tilt flexibility and twist flexibility per DNA sequence.

### Information content analysis

Information content per DNA binding site, Iseq ^48, 49^, was calculated separately for the whole set of REs as well as for the flexible and the rigid functional outcome categories, on the mononucleotide level, using the following equations:

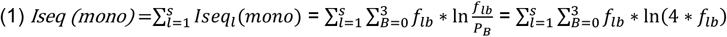

Where *Iseq_l_* (mono) is the mononucleotide information content per position, *f_lB_* is the frequency of occurrence of base *B* in position *l* in the population of all possible binding sites, and *P_B_* of base *B* is the frequency in the whole genome, which is usually taken to be 0.25 for each base. To correct for small sample size, we replaced equation (1) with the following equation, shown to be the best estimate for the sequence information when *f_lB_* is unknown precisely ^49^:

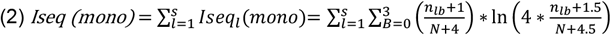

Where *N* is the sample size, and *n_lb_* is the number of occurrences of base-pair *B*(*B* = 0, 1, 2, 3) at position *l*. The expected variance in assigning *f_lB_* values was calculated from ^48^:

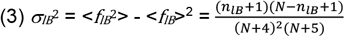

### Other computational and statistical tests

The values for the deformability (V(B) in units of °^3^Å^3^) of each DNA base-pair step were calculated as previously described ^34^, using the values of ^36^. Statistically significant differences in DNA deformability between p53-dependent functional-outcome groups, as well as analysis of the differences in transactivation level of the studied REs, were tested using one-way analysis of variance (ANOVA). We used the Dunn method for joint ranking as a post-hoc test, to determine the groups that are significantly different from each other by their deformability, since this method is adjusted for multiple non-parametric comparisons. We used student’s t-test as a post-hoc test for the differences between transactivation levels. All *P* values used in this study are two-sided. *Z* (Table S3) denotes the standardized test statistic, which has an asymptotic standard normal distribution under the null hypothesis of no difference in means. In testing the strength of the relationship between transactivation assays from reporter gene assays (RGA) and deformability (V(B)), we used Pearson correlation, using all data points from the biologically independent RGA experiments. All analyzes were conducted using JMP^®^, Version 16 (SAS Institute Inc., Cary, NC, 2021).

## CRediT authorship contribution statement

**Jessy Safieh:** Methodology, Software, Validation, Formal analysis, Investigation. **Ariel Chazan:** Methodology, Validation, Investigation. **Pratik Vyas:** Investigation. **Hanna Saleem:** Resources. **Yael Danin-Poleg:** Formal analysis; Writing - Review & Editing. **Dina Ron:** Supervision, Resources, Writing - Review & Editing. **Tali E. Haran:** Conceptualization, Supervision, Funding acquisition, Formal analysis, Data Curation, Writing - Original Draft, Writing - Review & Editing

## Data availability

Data will be made available on request.

## Acknowledgments

We thank Varda Rotter (Weizmann Institute of Science, Rehovot, Israel) for the p53 expression plasmid, pC53-SN3. We thank Alberto Inga (CIBIO, Trento, Italy) for stimulating discussions of the manuscript. We thank the biomedical core facility (BCF, Technion) for Sanger sequencing. This work was supported by the Israel Science Foundation (Grant #1517/14 to T.E.H.).

