## Supplementary Information for "Dynamics of p53 DNA binding sites contributes to functional selectivity of p53-driven gene expression"

**Faculty of Biology, Technion, Technion City, Haifa, Israel**

[1] present address: Department of Biomolecular Sciences, Weizmann Institute of Science, Rehovot, Israel

*To whom correspondence may be addressed.

**Table S2.** Average deformability values (V(B)$^{\circ 3}$Å$^{3}$) grouped by functional outcome of gene activation by p53. Listed groups are those with at least five genes.

| Functional outcome group | Gene symbols | Average V(B) per group ($^{\circ 3}$Å$^{3}$) ± SEM |
|---|---|---|
| Cell adhesion/anti-cell-motility | BRMS1L, CD82 KAI1, CDH10, CDH8, CLCA2, CLDN1, COL7A1, CSTA, DMTN, GALNT11, ICAM4, LPXN, PTPRU, SUSD6, WDR63 | 2.21 ± 0.03 |
| Cell growth | BTG4, GNAI1, LIMK2, SERTAD1, TEX14 | 2.30 ± 0.04 |
| Cell signaling | ADRB2, CAVIN2, CSNK1G1, FGF2, GDF15, GNA14, GPC1, HHAT, MGRN1, PDE2A, PDE4C, PHLDA3, PLK2, RRAD, S100A2, TAFA-2, TGFA, TNFAIP8, TRAF4 | 2.26 ± 0.04 |
| Cellular stress response | ALOX5, CPEB4, HMOX1, HSPA4L, ISCU, PPP1R14C, RFX7, SESN1, SESN2, SLC40A1 | 1.89 ± 0.02 |
| Cytoskeleton/cell motility | ACTA2, CALD1, CDC42BPG, CDC42EP3, CEP85L, COBLL1, CRPPA, CYRIA, EphA2, MMP2, NEFL, NUAK2, PARD6G, PLXNB1, PLXNB2, PSTPIP2, RAP2B, RHOC, SCRIB, VWCE | 1.99 ± 0.03 |
| Development | ADGRB2, ANK1, ASTN2, COL18A1, CPE NF-α1, ELFN1, ELFN2, EPS8L2, FANK1, KCNJ12, KIF5C, SULF2 | 2.52 ± 0.03 |
| DNA repair | ASCC3, PCNA, POLH, REV3L, RRM2B | 2.25 ± 0.08 |
| Early DDR steps | BTG2, CCNG1, CDKN1A, IER3, MCC, RPS27L, SFN, TP53I3 | 2.41 ± 0.04 |
| Energy homeostasis | CPE, GLS2, PANK1, PTPRE, TRIM55 | 2.56 ± 0.06 |
| Extrinsic apoptosis | CDIP1, FAS, RASSF5, TNFRSF10A, TNFRSF10B, ZC3HAV1 | 2.28 ± 0.04 |
| Innate immunity | APOBEC3C, APOBEC3H, CASP1, DTX4, GSDME, HERC5, MARCH8, PADI4 PAD4, PLEKHF1, RNASE7, RPS19, SP110, SPATA18, TLR3, TRIM22, TRIM26, TRIM35, TRIM5 | 2.45 ± 0.05 |
| Intrinsic apoptosis | ACER2, BAX, BBC3, BCL2L14, LAPTM5, PERP, PMAIP1, TP53AIP1, TRIML2 | 1.96 ± 0.04 |
| Ion homeostasis | SCN2A, SCN3B, SCN4B, SLC12A4, SLC25A45, SLC30A1, SLC9A1, TRPM1 | 2.26 ± 0.04 |
| Lipid homeostasis | CES2, CROT, DHRS3, FDXR, GPX1, IP6K2, METTL7A, PRKAB1 | 2.32 ± 0.04 |
| Membrane trafficking | ABCA12, AP4E1, LRPAP1, PGAP1, SORL1, SYTL1, TEPSIN, TMEM30A, EPN3 | 2.19 ± 0.05 |
| p53 negative regulation | MDM2, PPM1D, PRDM1, PTP4A1, XPC | 1.89 ± 0.06 |
| Stem cells & differentiation | ALDH1A3, ARHGEF3, CSF1, EDA2R, EFNB1, FAM210B, FBXW7, GRHL3, LIF, MSX2, NHLH2, SLC4A11, TMEM64, UBASH3B, ZFPM1 | 2.23 ± 0.03 |
| Survival | BTBD10, CFLAR, DYRK3, GDNF, GPR87, NGF, TMBIM1, TNFRSF10C, TNFRSF10D, TRIAP1 | 2.18 ± 0.05 |

**Table S3.** Significant differences between deformability values (V(B)°³Å³) grouped by functional outcome of gene activation by p53.

| First outcome group | to second outcome group | Score mean difference [a] | Std error difference [b] | *Z* [c] | *P*-Value |
|---|---|---|---|---|---|
| Development | Cellular stress response | 147.36 | 23.08 | 6.39 | <.0001 |
| Development | Cytoskeleton/cell motility | 127.34 | 19.99 | 6.37 | <.0001 |
| Development | Intrinsic apoptosis | -135.62 | 24.33 | -5.57 | <.0001 |
| Development | p53 negative regulation | -147.33 | 29.19 | -5.05 | <.0001 |
| Early DDR steps | Cellular stress response | 126.57 | 25.15 | 5.03 | <.0001 |
| Early DDR steps | Cytoskeleton/cell motility | 106.55 | 22.35 | 4.77 | 0.0003 |
| Early DDR steps | Intrinsic apoptosis | -114.83 | 26.30 | -4.37 | 0.0019 |
| Early DDR steps | p53 negative regulation | -126.54 | 30.86 | -4.10 | 0.0063 |
| Energy homeostasis | Cellular stress response | 150.14 | 29.19 | 5.14 | <.0001 |
| Energy homeostasis | Cytoskeleton/cell motility | 130.11 | 26.81 | 4.85 | 0.0002 |
| Energy homeostasis | Intrinsic apoptosis | -138.39 | 30.19 | -4.58 | 0.0007 |
| Energy homeostasis | p53 negative regulation | -150.10 | 34.23 | -4.38 | 0.0018 |
| Innate immunity | Cellular stress response | 124.39 | 20.71 | 6.01 | <.0001 |
| Innate immunity | Cytoskeleton/cell motility | 104.36 | 17.20 | 6.07 | <.0001 |
| Innate immunity | Intrinsic apoptosis | -112.64 | 22.10 | -5.10 | <.0001 |
| Innate immunity | p53 negative regulation | -124.35 | 27.36 | -4.54 | 0.0008 |
| Lipid homeostasis | Cellular stress response | 102.57 | 25.15 | 4.08 | 0.0069 |
| Cell signaling | Cellular stress response | -83.08 | 20.506 | -4.05 | 0.0078 |

[a] Score mean difference: The mean of the rank score of the deformability values in the first outcome group minus the mean of the rank scores of the deformability values in the second outcome group.

[b] Std error difference. The standard error of the score mean difference.

[c] Z: The standardized test statistic.

**Table S4:** Primers used for incorporation of p53 REs into pCluc Mini-TK plasmids

| **Gene symbol** | **Primer direction** | **Sequence (5’->3’)** |
|---|---|---|
| RRM2B | Forward | AGGCATGTCTTCCAAGGATCCTTCGCATATTAAGGTGAC |
| | Reverse | GGGCATGTCAGGACAGAATTCCAAGATCTCCCGATCCG |
| CCNG1 | Forward | AGGCTAGTCCGAGGCGGATCCTTCGCATATTAAGGTGAC |
| | Reverse | GGGCTTGTGCTCACAGAATTCCAAGATCTCCCGATCCG |
| p21-’5 | Forward | CAACATGTTGAGCTCGGATCCTTCGCATATTAAGGTGAC |
| | Reverse | GGACATGTTCCTGACGAATTCCAAGATCTCCCGATCCG |
| PMAIP1 | Forward | GGGCAGGTCGCGCTCGGATCCTTCGCATATTAAGGTGAC |
| | Reverse | GGACACGCTCCCGACGAATTCCAAGATCTCCCGATCCG |
| TP53AIP1 | Forward | GGGCTTGTCGAGATGGGATCCTTCGCATATTAAGGTGAC |
| | Reverse | GGGCAAGAGAGGAGGGAATTCCAAGATCTCCCGATCCG |

**Table S5:** Sequences used in the cyclization kinetics assays

| **Functional outcome** | **Gene symbol** | **Test sequences in cyclization kinetics assays [a]** |
|---|---|---|
| DNA repair | RRM2B | tgtccTGACATGCCCAGGCATGTCTtccaa |
| Cell cycle arrest | CCNG1 | tgtgaGCACAAGCCCAGGCTAGTCCgaggc |
| Cell cycle arrest | p21-5` | gtcagGAACATGTCCCAACATGTTGagctc |
| Apoptosis | PMAIP1 | gtcggGAGCGTGTCCGGGCAGGTCGcgctc |
| Apoptosis | TP53AIP1 | cctccTCTCTTGCCCGGGCTTGTCGagatg |

[a] Capital letters are the sequences of natural p53 REs. Lower letters are the natural flanking sequences

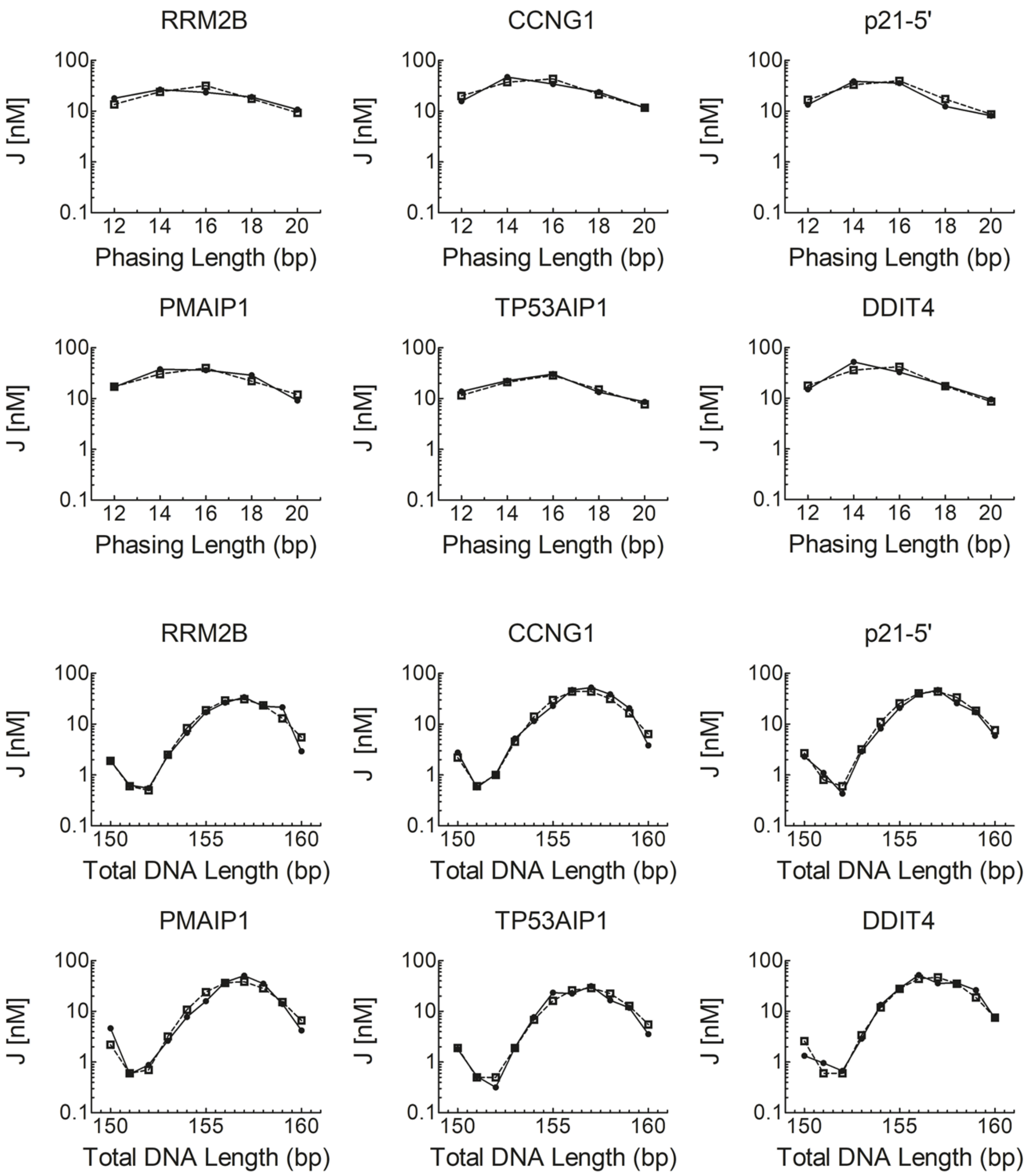

**Figure S1.** Cyclization kinetics of p53 REs. The J-factors for the DNA constructs as a function of either the phasing length (top two rows), or the total DNA length (bottom two rows). The solid lines are the experimental curves and the dashed lines are the curves from simulating the data.

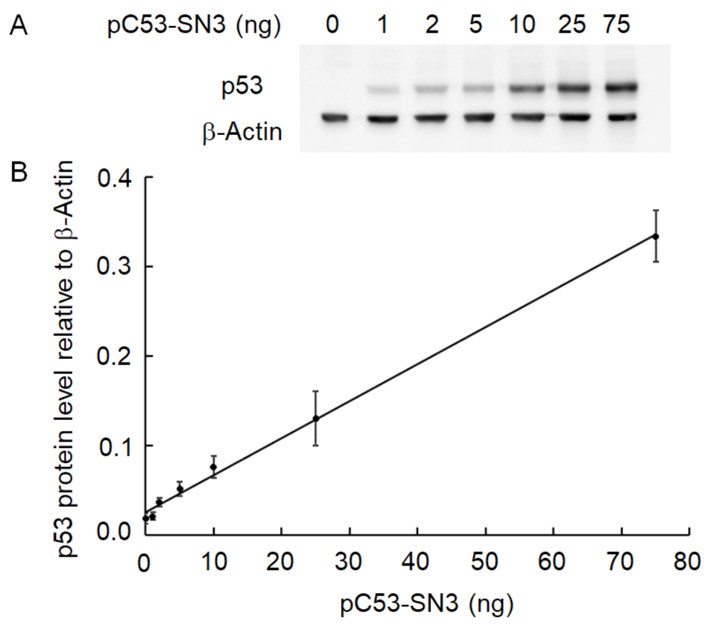

**Figure S2**: Validation of ectopic p53 expression level in H1299 cells. (A) H1299 cells were transfected with increasing amount of p53 expression vector. After 48 h, 10 µg of whole cells lysate was analyzed by immunoblotting with an antibody against p53. β-Actin serves as control. (B) Correlation between pC53-SN3 plasmid amount at the transfection against p53 protein level relative to β-actin, derived from (A). p53 protein level was estimated by normalizing the band intensity of p53 protein to the band intensity of β-Actin from the same sample. There is a linear relationship between the amounts of p53 expressing plasmid used in the transfection and the observed p53 protein levels (R= 0.998). The results are averages of four independent experiments.

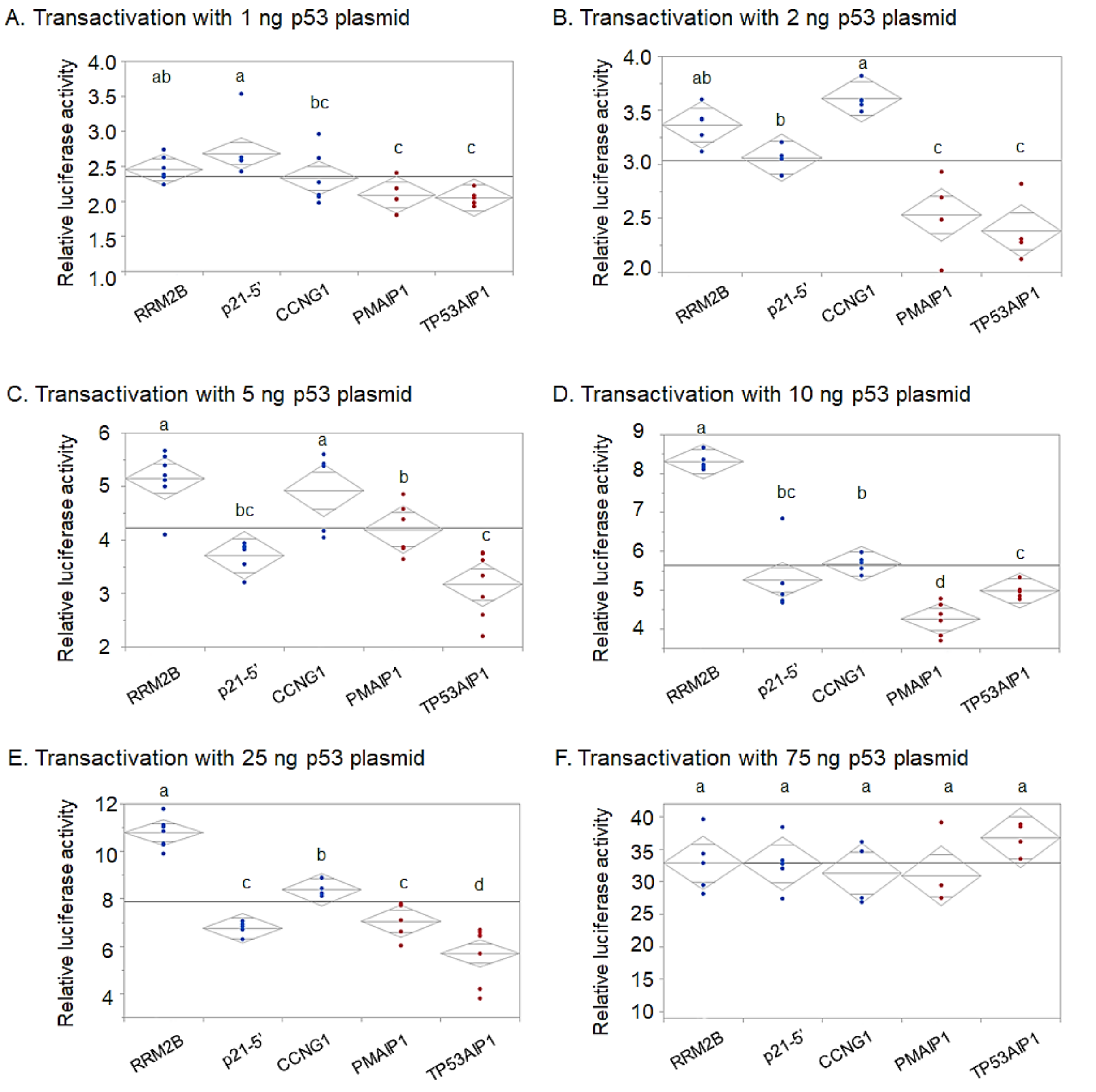

**Figure S3**. Variance analysis of transactivation from five p53 REs as a function of p53 plasmid levels tested by reporter-gene assay. The dots are individual independent assays with each RE. The line across each diamond represents the group mean. The vertical span of each diamond represents the 95% confidence interval (CI) for each group. Overlap marks $((\sqrt{2}\text{CI})/2)$ are drawn above and below the group mean. Means comparison was carried out by Student's t-test, at $\alpha = 0.05$ level. REs not connected by the same letter have significantly different transactivation levels.

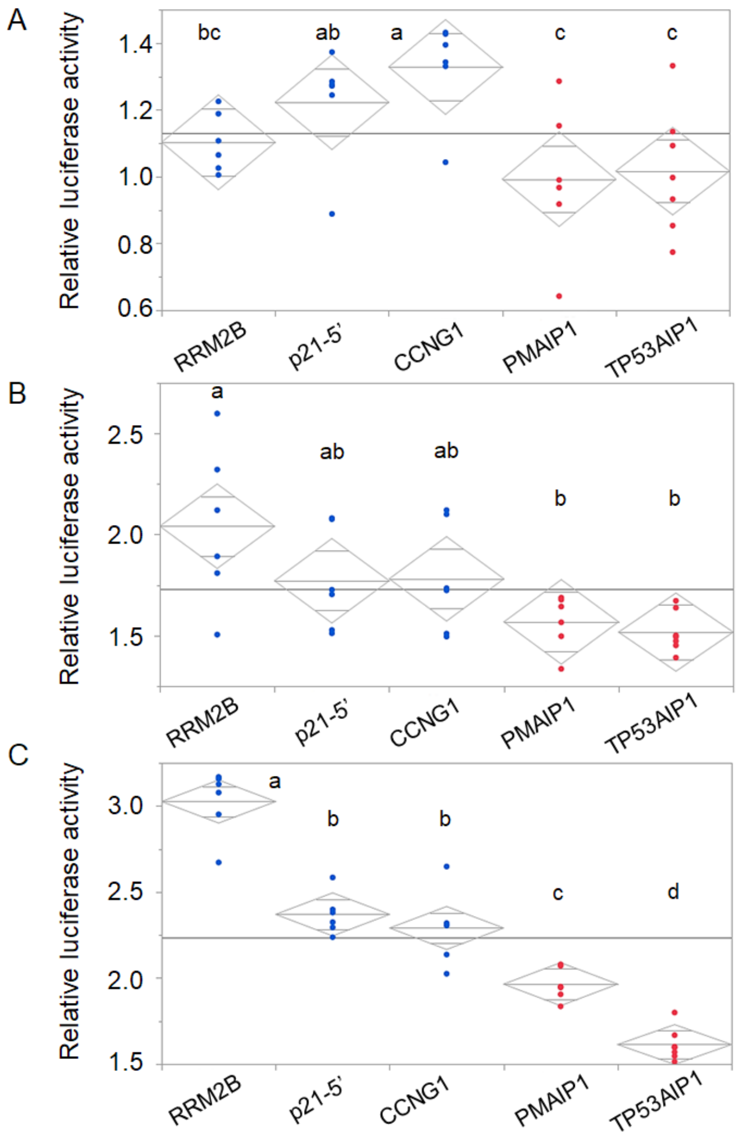

**Figure S4.** Variance analysis of transactivation from five p53 RE as a function of time, tested by reporter-gene assay and 25 ng p53 plasmid. The line across each diamond represents the group mean. The vertical span of each diamond represents the 95% confidence interval for each group. Overlap marks $((\sqrt{2}CI)/2)$ are drawn above and below the group mean. Means comparison was by Student's t-test, at $\alpha = 0.05$ level. Levels not connected by the same letter are significantly different. (A) 12 h post-transfection, (B) 18 h post-transfection, (C) 24 h post-transfection.
